# Brisk: Exact resource-efficient dictionary for *k*-mers

**DOI:** 10.1101/2024.11.26.625346

**Authors:** Caleb Smith, Igor Martayan, Antoine Limasset, Yoann Dufresne

## Abstract

The rapid advancements in DNA sequencing technology have led to an unprecedented increase in the generation of genomic datasets, with modern sequencers now capable of producing up to ten terabases per run. However, the effective indexing and analysis of this vast amount of data pose significant challenges to the scientific community. K-mer indexing has proven crucial in managing extensive datasets across a wide range of applications, including alignment, compression, dataset comparison, error correction, assembly, and quantification. As a result, developing efficient and scalable *k*-mer indexing methods has become an increasingly important area of research. Despite the progress made, current state-of-the-art indexing structures are predominantly static, necessitating resource-intensive index reconstruction when integrating new data. Recently, the need for dynamic indexing structures has been recognized. However, many proposed solutions are only pseudo-dynamic, requiring substantial updates to justify the costs of adding new datasets. In practice, applications often rely on standard hash tables to associate data with their *k*-mers, leading to high *k*-mer encoding rates exceeding 64 bits per *k*-mer. In this work, we introduce Brisk, a drop-in replacement for most *k*-mer dictionary applications. This novel hashmap-like data structure provides high throughput while significantly reducing memory usage compared to existing dynamic associative indexes, particularly for large *k*-mer sizes. Brisk achieves this by leveraging hierarchical minimizer indexing and memory-efficient super-*k*-mer representation. We also introduce novel techniques for efficiently probing *k*-mers within a set of super-*k*-mers and managing duplicated minimizers. We believe that the methodologies developed in this work represent a significant advancement in the creation of efficient and scalable *k*-mer dictionaries, greatly facilitating their routine use in genomic data analysis.

## 1 INTRODUCTION

Handling sequencing and genomic data is among the most memory-intensive tasks routinely performed in computational biology (Stephens et al., 2015; Cremin et al., 2022). Superlatives abound in this field, with the mistletoe genome nearing 100 gigabases (GCA 963277665.1), recent sequencers capable of outputting 16 terabases per run, and future developments promising even more orders of magnitude improvement^1^. Additionally, genome databases like GenBank are rapidly expanding, now encompassing over 29 terabases of data^2^. Managing these massive datasets demands significant computational resources, driving the development of high-performance tools for commonly performed tasks such as alignment, assembly, and genotyping.

Since the advent of BLAST, a fundamental need has emerged to index *k*-mers and associate information with them (Jenike et al., 2024; Marchet, 2024a,b). While *k*-mers offer several advantages, including efficient filtering, sensitivity/specificity trade-offs, and convenient representation as integers, they exacerbate memory usage challenges. A text of *N* bases generates *𝒪*(*N*) *k*-mers (assuming no repeats), which represent *𝒪*(*N·k*) DNA bases. With commonly used *k*-mer sizes ranging from 17 to 31, *k*-mers are often represented by 64-bit integers, leading to an 8-byte per *k*-mer footprint. Consequently, handling gigabase-sized documents, which are already extremely memory-intensive, typically requires workstation-level resources.

Surprisingly, most tools still utilize generic dictionary/hashmap structures for *k*-mer indexing, despite their high memory costs, as the 8-byte per *k*-mer footprint is further multiplied due to the load factor. A viable solution to mitigate this memory usage is to avoid indexing all *k*-mers, instead using minimizers to ensure the scalability of tools (Li, 2018; Benoit et al., 2024; Rouzé et al., 2023; Agret et al., 2022). Recently, indexes capable of associating *k*-mers with additional information have been proposed. These indexes generally fall into two categories: full-text indexes based on the Burrows-Wheeler Transform (BWT) (e.g., BOSS (Bowe et al., 2012), Themisto (Alanko et al., 2023)) or Minimal Perfect Hash Functions (MPHF) (e.g., BBHash (Limasset et al., 2017), PTHash (Pibiri and Trani, 2021), Pufferfish (Almodaresi et al., 2018), Blight (Marchet et al., 2021), SShash (Pibiri, 2022)).

While these structures can achieve very low memory footprints (below 8 bits per *k*-mer) and high throughput, they present two major caveats. To reduce the memory footprint, these methods rely on assembling *k*-mers into sequences that can contain them in a memory-efficient manner (Rahman and Medevedev, 2021). Therefore, they depend on such spectrum-preserving string sets (SPSS) structures to construct their index, meaning that *k*-mers must be “assembled” before the indexing step. Despite significant progress in this area, this remains a resource-intensive process (e.g. Bcalm2 (Chikhi et al., 2016), Cuttlefish (Khan and Patro, 2021), ggcat (Cracco and Tomescu, 2023)).

In a subsequent step, the index structures, either MPHF or BWT-based, must be built. Consequently, while these indexes are resource-efficient, they are intrinsically static and require substantial resources to construct. These efficient yet costly-to-construct indexes are particularly valuable in scenarios where the index is reused multiple times for a large number of queries. However, in most applications, the number of queries is relatively low, and the construction cost is not justified. As a result, the construction of such indexes is often avoided, as it would be the time bottleneck of the application.

Another significant issue with static indexes is the need for complete rebuilding when incorporating new datasets. If updates are frequent, the building costs become overwhelming. Some semi-dynamic approaches have been proposed, using temporary buffers and partitions to delay reconstruction as much as possible (Alanko et al., 2021; Almodaresi et al., 2022). However, these approaches introduce latency, as some insertions trigger costly rebuilding operations, leading to periods when the index is unavailable. Moreover, these methods are typically unsuitable for streaming applications, as they require multiple passes over the dataset.

These approaches also assume that the set of *k*-mers to be indexed is known in advance. In such cases, decisions can be made about whether to index a *k*-mer based on the current state of the index, for example, avoiding indexing a *k*-mer if a very similar one is already present. However, these methods cannot be used in streaming, and, more critically, the indexing phase cannot be leveraged to associate data with the *k*-mer as it is inserted, unlike a dictionary, which can associate data with the *k*-mer immediately upon insertion. Consequently, most applications require an index that can be quickly constructed in streaming with *k*-mer-level dynamism.

Such indexes are quite rare. Jellyfish (Marcais and Kingsford, 2012), which is very close to a classical hash table, provides good performance by being lock-free. CBL (Martayan et al., 2024a) improves on this scheme by exploiting the locality of successive *k*-mers. Both of these structures achieve very fast insertion and query times; however, since they do not rely on SPSS, they exhibit memory usage in *𝒪*(*N*.*K*). Bifrost (Holley and Melsted, 2020) is the only dynamic tool that actually relies on SPSS (unitigs in this case) to reduce memory usage, but it is almost an order of magnitude slower than a hash table.

In this work, we aim to propose a drop-in replacement for *k*-mer dictionaries for many use cases, offering fast query/insertions while being orders of magnitude more memory-efficient than other dynamic methods, and providing *k*-mer-level dynamism.

## 2 METHODS

### 2.1 Outline

In order to minimize the memory footprint of our dictionary, we aim to represent *k*-mers as SPSS. However, most SPSS representations such as simplitigs (Rahman and Medevedev, 2021), matchtigs (Schmidt et al., 2023), eulertigs (Schmidt and Alanko, 2023), and masked superstrings (Sladky et al., 2023) are defined globally from a complete *k*-mer set. Updating such SPSS structures remains an open problem, making their use in dynamic structures even more challenging (Hannoush et al., 2024). Interestingly, one of the first SPSS used was super-*k*-mers, which are successive *k*-mers sharing a minimizer (Deorowicz et al., 2015). While this structure is more memory-intensive than other SPSS types, it remains lighter than plain *k*-mer representation by several folds.

The main advantage of super-*k*-mers is their locality; they can be built incrementally while parsing a sequence with minimal overhead, making them well-suited for dynamic usage. Another significant advantage of super-*k*-mers over more space-efficient SPSS structures is that we can determine, without any additional context, to which minimizer a *k*-mer is associated, allowing us to use this minimizer for clustering *k*-mers (Marchet et al., 2020; Marchet and Limasset, 2023).

By grouping all super-*k*-mers associated with the same minimizer, we can efficiently determine in which bucket a *k*-mer belongs, enabling searches within a smaller substructure. Furthermore, since all super-*k*-mers in a bucket contain the same minimizer, we can skip the encoding of the minimizer in the super-*k*-mers, only encoding the prefixes and suffixes before and after the minimizer (Rouzé et al., 2023, Partitioned sketches).

However, within a set of *S* super-*k*-mers sharing a minimizer (often referred to as a bucket), checking the presence of a given *k*-mer requires *𝒪*( |*S*|) comparisons, which can be costly. To complicate matters, buckets are known to have skewed size distributions, as lower minimizers tend to gather significantly more *k*-mers than others leading to expensive search for a large number of *k*-mers (Chikhi et al., 2014; Deorowicz et al., 2015; Marchet and Limasset, 2023). Another challenge is that a *k*-mer may have multiple occurrences of its minimizers, potentially causing issues if a *k*-mer/super-*k*-mer encoding depends on which minimizer is selected during parsing. Therefore, we need a deterministic method to select a “canonical” minimizer when multiple occurrences arise.

Our practical dictionary implementation addresses these issues based on four key contributions, each discussed in the following sections:

- Indexing a low amount of memory-efficient super-*k*-mers using state-of-the-art minimizer scheme.
- Leveraging large minimizers through lazy encoding, enabled by careful handling of duplicated minimizers.
- Sorting super-*k*-mers to accelerate fast probing.
- Using superbuckets to achieve uniform bucket distribution.

### 2.2 Indexing Super-*k*-mers

Since their introduction, minimizers have been widely used across a broad range of applications (Deorowicz et al., 2015; Li, 2018; Marchet et al., 2020; Ekim et al., 2023; Pibiri et al., 2023; Marchet and Limasset, 2023). A very powerful property of minimizers is that while every *k*-mer has a minimizer of size *m*, a trivial lower bound for the fraction of *m*-mers being minimizers, known as density, is 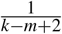 A simple scheme based on hashed *m*-mers achieves a density of 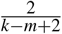 (Schleimer et al., 2003). In practice, this means that one can select a set of minimizers that is an order of magnitude smaller than the *k*-mer set. As a result, the corresponding super-*k*-mers can group numerous *k*-mers together. For random minimizer density, we can theoretically expect (*k− m* + 2)*/*2 *k*-mers per super-*k*-mer, assuming they are extracted from a long sequence with appropriate parameters. In Figure 1a, we show the mean super-*k*-mer size that can be obtained for standard values of *k* and *m*, and observe that practical results closely match this approximation. This leads to an expected memory footprint of 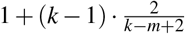nucleotides per *k*-mer (Martayan et al., 2024b, Theorem 1), i.e. 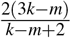 bits per *k*-mer. In practice, since super-*k*-mers have distinct sizes, an additional cost of log(*k – m + 2)* bits must be added to encode the super-*k*-mer size. A simpler practical alternative is to allocate the maximum size for every super-*k*-mer. If we neglect *k*-mer with duplicated minimizer, the maximal size of a super-*k*-mer correspond to the case where the first *k*-mer has its minimizer at the rightmost position and the last *k*-mer has its minimizer at the leftmost position. These represent 2*k − m* bases encoding 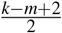 *k*-mers and provide a bits-per-*k*-mer ratio of 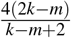. While this representation is more costly, it offers very convenient properties such as constant access to elements in the list and efficient preallocation, enabling sublinear search within the list, which we aim to implement in this work.

**Figure 1.**
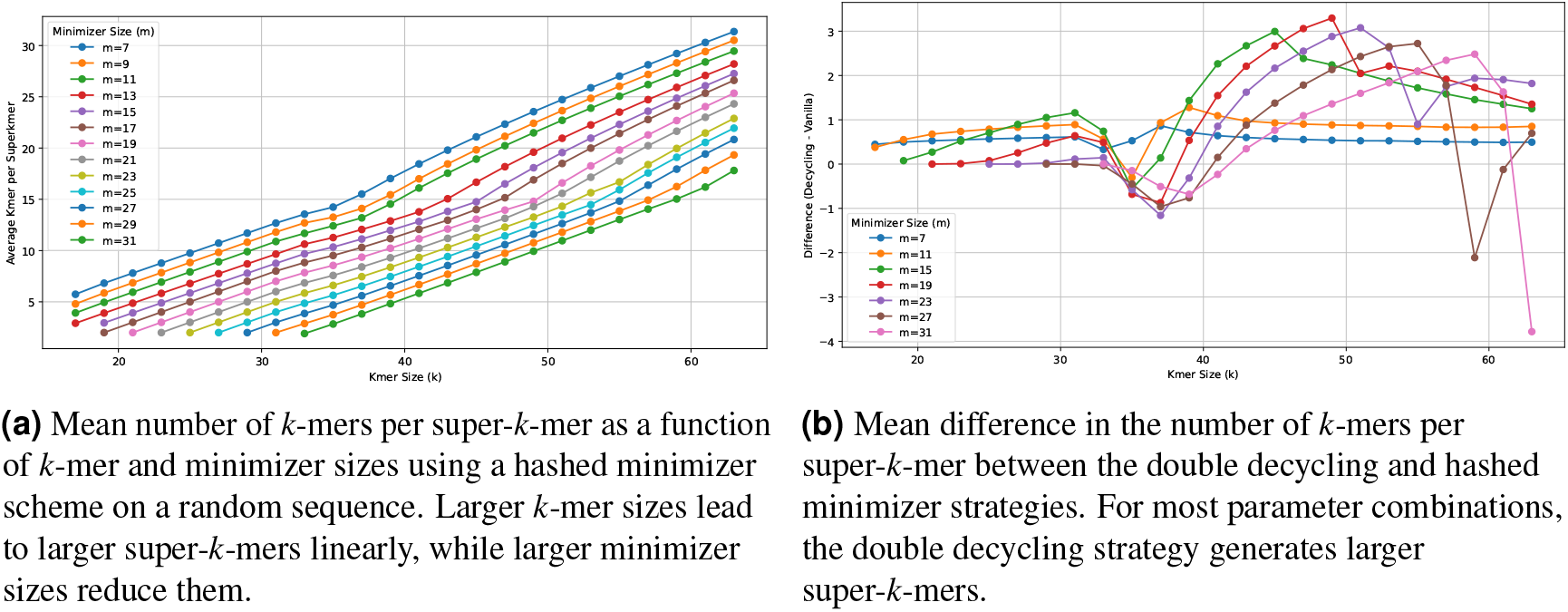
Analysis of Super-*k*-mers Statistics and Strategy Differences. (a) Shows the mean number of *k*-mers per super-*k*-mer based on varying *k*-mer and minimizer sizes. (b) Illustrates the mean difference in *k*-mers per super-*k*-mer between double decycling and hashed minimizer strategies.

This memory cost assumes the use of hashed minimizers that provide a density of 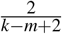. Being able to select fewer minimizers would result in fewer, larger super-*k*-mers and a reduced memory cost. Several works have aimed to further improve minimizer density, such as miniception (Zheng et al., 2020), decycling sets (Pellow et al., 2023), and mod-minimizers (Groot Koerkamp and Pibiri, 2024). In this work, we pragmatically chose to use the decycling set scheme (Pellow et al., 2023) as an improved minimizer scheme for its computational efficiency and practical improvement in super-*k*-mers length on our parameter range. In Figure 1, we display the improvement in mean super-*k*-mer length using the double decycling set compared to the random minimizer scheme. Since these minimizers can be selected in streaming with minimal computational overhead, we use them to benefit from a reduced number of minimizers to index, resulting in larger super-*k*-mers and an improved bits-per-*k*-mer ratio.

### 2.3 Lazy Encoding and Multiple Minimizers

When encoding all super-*k*-mers associated with a given minimizer, we can exploit the fact that the minimizer sequence is present in each super-*k*-mer. Moreover, in a maximal super-*k*-mer, the position of the minimizer is known, so it can be skipped entirely. This results in a bits-per-*k*-mer ratio of 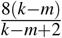 per super-*k*-mer if we neglect the cost of storing the minimizer, as it is encoded once per bucket. For practical values of *k* and *m*, 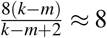, which is roughly one byte per *k*-mer.

Although this lazy encoding strategy reduces memory usage, it requires each *k*-mer’s minimizer position and strand within its sequence to be determined consistently. While this is not an issue in most cases where the minimal *m*-mer sequence occurs only once within its *k*-mer, situations with multiple occurrences are often overlooked and labeled as edge cases. Additionally, some studies disregard the reverse complement aspect of DNA sequences, which can have a significant impact in practice (Marçais et al., 2024).

Selecting a minimizer position also implies choosing the strand of the *k*-mer, as a *k*-mer and its reverse complement can have the same, canonical, minimizer at different positions, leading to ambiguity in how the *k*-mer is stored. In the following, we define a complete and deterministic method to select a given occurrence of the minimal *m*-mer across a *k*-mer sequence or its reverse complement.

To determine the minimizer position and strand in these cases, we check both the *k*-mer sequence and its reverse complement and apply three selection rules to ensure consistency:

1. Canonical minimizer
2. Leftmost canonical minimizer
3. Leftmost canonical minimizer from canonical *k*-mer

In most cases, as shown in Figure 2A, the minimizer appears once in the *k*-mer but also appear in its reverse complement. The first rule is to consider only canonical *m*-mers (*m*-mers smaller than their reverse complements) as potential minimizers. This fix the *k*-mer strand as only one strand contain the canonical minimizer (Figure 2A).

**Figure 2.**
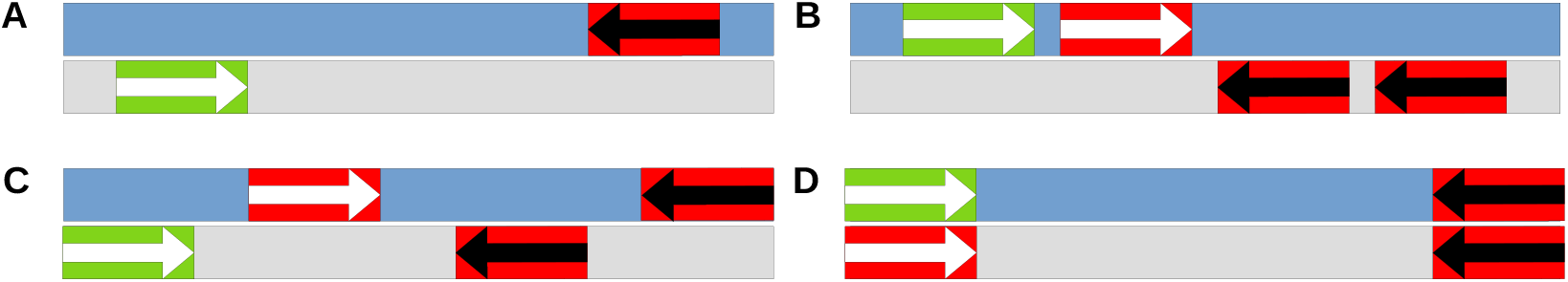
This figure illustrates four cases of canonical minimizer selection within a *k*-mer. The blue bars represents the canonical *k*-mers, while the gray bars represents the complementary *k*-mers. The occurrences of a *k*-mer minimizer are represented by arrows: white arrows for minimizers whose *m*-mer is canonical and black arrows for non-canonical ones. The minimizer occurrence highlighted in green is the one selected, while the red ones are not selected. Case A presents the most common scenario, where there is no duplicated minimizer; the one with the canonical sequence is selected following rule 1. Case B presents a duplicated minimizer where both occurrences appear canonical on the same strand, while Case C presents canonical occurrences on both strands. In either case, following rule 2, the canonical occurrence at the leftmost position within its own strand is selected. Case D is the rare case where the leftmost canonical occurrences of each strand are at the same position. In such cases, following rule 3, we favor the one in the canonical *k*-mer.

However, in a minority of cases, the canonical minimizer occurs more than once within a *k*-mer. This duplication can occur either on the same strand (Figure 2B) or across both the canonical and complementary *k*-mer (Figure 2C). To handle this, we apply the second rule: always select the leftmost occurrence of the canonical minimizer among the *k*-mer and its reverse complement. In Figure 2B, where both occurrences are on the same strand, the leftmost minimizer on this strand is chosen. In Figure 2C, the minimizer in the reverse complement *k*-mer is selected because its position is lower than the corresponding occurrence in the canonical strand.

A rare edge case arises when two leftmost canonical minimizers exist simultaneously: one in the canonical *k*-mer and one in its reverse complement. These “mirror” cases are resolved by selecting the minimizer in the canonical *k*-mer as depicted in Figure 2D. Note that the parity of *k* does not complicate this process. If *k* is odd, palindromic *k*-mers are impossible, so one strand is inherently superior. For even *k*, palindromic *k*-mers are possible, but both strands are equally canonical, ensuring no ambiguity.

By following these rules, every *k*-mer is deterministically associated with a strand and a minimizer position.

### 2.4 Super-*k*-mer probing

Super-*k*-mers are widely used as memory efficient *k*-mers representation. A reason for this, beside the ones previously mentioned, is that a given *k*-mer has at most one possible potential position in a given super-*k*-mer (zero if they have different minimizer) obtained from their relative minimizer position. However, given a *k*-mer, its corresponding bucket containing all sharing its minimizer with it can be extremely large and performing all comparison computationally expensive, so the objective is to accelerate such search. A set of *k*-mers can be placed in a hashmap like data structure to get practical constant query time but hashing super-*k*-mers would not allow us to perform *k*-mer inclusion queries. Similarly, probing a *k*-mer in a sorted *k*-mer list can be performed efficiently sublinear time using binary search but sublinear probing in a super-*k*-mer list is not trivial as sorted super-*k*-mers are irrelevant to the *k*-mer location in the list.

To improve from naive linear probing we introduce a novel super-*k*-mers representation that we call interleaved super-*k*-mers and show that searching in a sorted list of interleaved super-*k*-mers can lead to sublinear probing.

Intuitively sorting super-*k*-mers lexicographically is not a good idea because it gives most importance to the first bases. This seems irrelevant for at least two reasons. First the left bases have crucial impact while right are mostly irrelevant despite having symmetrical properties. Secondly, the leftmost (resp rightmost) bases are the less important in the sense is that are shared by less *k*-mer than “central” bases that surround the minimizer. If the minimizer is shared by all *k*-mer, the leftmost (resp rightmost) base is exclusive to a given *k*-mer, the next base is shared by two *k*-mer and so on. Inversely the base next to the minimizer are shared by every *k*-mers but one, the next by every *k*-mer but two and so on. Based on this observation we propose a base reordering that prioritize the most shared bases. The interleaved transformation of a super-*k*-mer starts by its minimizer sequence and is followed by the next base at its right then the base at its left and repeating this process iteratively alternating left and right bases. When no bases are available, an N character is used to encode this absence. Note that in practice, we do not write the Ns explicitly, but we infer the positions that can be ignored using the length and the offset of the super-*k*-mer. Some example of this transformation are showed in Figure 3.

**Figure 3.**
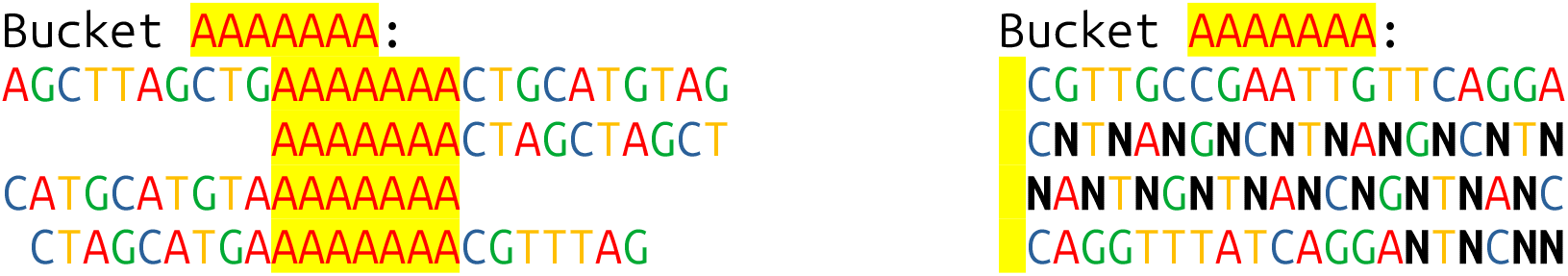
Toy example of the AAAAAAA bucket associated to four super-*k*-mers turned into their interleaved representation.

As premised the bases are ordered according to how likely they are to be shared by numerous *k*-mers. Such transformation can also be applied on a *k*-mer as it is a super-*k*-mer itself. An interesting property of this transform is that a *k*-mer is included in a super-*k*-mer if and only if the interleaved *k*-mer is a prefix of the super-*k*-mer interleaved (considering that the N character match with any other). Note that this property works for any super-*k*-mer and not exclusively for the *k*-mer case. We also highlight that this property would hold for any deterministic order over the positions of a super-*k*-mer, but we chose the positions that are less likely to be an N. While this property does not grant anything in itself as it does not simplify the *k*-mer presence test, it allows the interleaved super-*k*-mers to be sorted and to perform a binary search. Several examples of binary search are displayed in Figure 4, we observe that very few steps are necessary to find a given location as every base divide the search space by four. However, when there is an N in the *k*-mer it could be present in a super-*k*-mer with any nucleotides at this position, so the search space is not reduced and the four possibilities have to be checked with the following bases. This mean that from the first N appearance the probing will alternate between a useful base that divide the search space and an exhaustive search impeding the search. If this happens at the end of the probing when the search space is small this hardly impact search time. However, if this happened in the first bases when the search space is very large the actual probing because closer to a linear probing. The specific worst case for our representation is when *k*-mers start or end with their minimizer as it generate N characters from the beginning of the interleaved. Such *k*-mers are the most difficult to probe as displayed in Figure 4. However all other cases benefit from exponential reduction of the search space from first bases.

**Figure 4.**
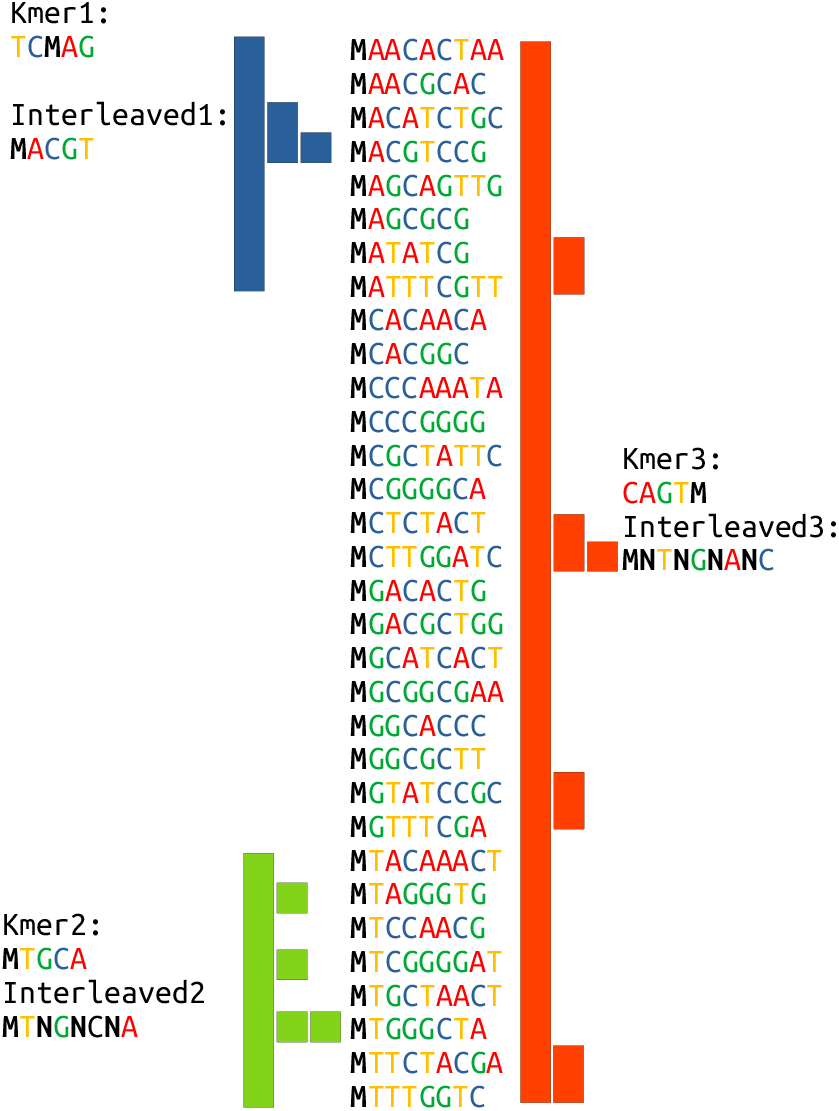
Examples of *k*-mer search in a bucket of 32 super-*k*-mers. M denotes the bucket minimizer. The colored bar indicates the successive possible search spaces after each step. *k*-mer 1 search represented by blue bars start with A, then C that reduces the search space and G that limits the search to one *k*-mer and end it. *k*-mer 2 search represented by green bars starts with T but the next char is a N so it not useful and the search space is unchanged, the next character is a G, that provides several smaller search space, once again a N char that cannot be used and finally the C limits the search to one *k*-mer and ends it. *k*-mer 3 search represented in red bars starts with a N so every position is a possible match, the second character T provides four distinct search space, the next char N does not help, the next char G limit the search to one position and end it.

### 2.5 Superbuckets

Since our approach indexes super-*k*-mer lists associated with a minimizer, which we call a bucket, the size of the chosen minimizer, *m*, is crucial. For a given *k*, a smaller minimizer can generate fewer but larger super-*k*-mers. However, since there are *𝒪*(4^*m*^) possible minimizers, a larger minimizer results in more, smaller buckets that are faster to probe. Therefore, the minimizer size acts as a time-memory trade-off. Although increasing *m* by two (to keep it odd) can slightly improve the mean number of *k*-mers per super-*k*-mer (see Figure 1a), it also multiplies the number of potential buckets by 16, significantly impacting the mean bucket size. Thus, the advantages of larger minimizers tend to outweigh those of smaller ones.

However, due to various factors—both biological (such as repeats or distribution biases) and computational (such as reverse complement considerations and the fact that minimizers are minimal over *k − m* + 1 other values)—minimizers are often poorly distributed, leading to two distinct problems: a large number of empty buckets and some very large buckets (Pibiri, 2022). To illustrate this phenomenon, we plot the bucket size distribution for the *C. elegans* reference genome compared to a random sequence of the same size in Figure 6. We observe that; for the three *m* values, no bucket from the random sequence contains 2^6^ or more elements, while buckets from *C. elegans* can contain more than 2^10^ elements. We also see that *C. elegans* generates more very small buckets than the random sequence. These observations highlight the skewed nature of genomic data that negatively impact on our scheme.

To address this issue, we introduce the concept of superbuckets, which merge multiple buckets into one in a uniform manner. Since the problem lies in the non-uniform distribution of minimizers, a simple solution is to use a hash function to achieve a uniform distribution. However, using a non-injective function would lose information and allow hash collisions for different minimizers, leading to false-positive matches. To prevent this, we use a bijective/invertible function that guarantees the original minimizer can be retrieved from its hashed value. We then use the hashed minimizer prefix to group multiple buckets together. As bijective hashing change the order without changing the distribution, buckets are uniformly grouped together, smoothing the difference in number of *k*-mers inside the super-buckets.

In practice, only 4^*b*^ (*b < m*) superbuckets are used, based on the first *b* nucleotides of the interleaved super-*k*-mers with hashed minimizers. Since the hashed minimizer bits are uniformly distributed, the number of buckets per superbucket is also uniform. Although the bucket sizes are not uniform, superbucket sizes are less skewed. Figure 6 shows the size distribution of regular buckets versus superbuckets with different *b* parameters: *m* = *b* + 2 results in an average of 16 buckets per superbucket, *m* = *b* + 4 results in 256 buckets per superbucket, and *m* = *b* + 6 results in 4096 buckets per superbucket. We observe that smaller *b*, relative to *m*, leads to fewer large buckets and more small ones, improving overall bucket size and consequently reducing probing time.

This optimization has minimal drawbacks, as minimizers (and therefore *k*-mers) can be reconstructed since an invertible hash is used, as the hashed minimizer value can be reconstructed from its superbucket information. One might argue that merging small buckets into larger superbuckets could hinder searching. However, because these superbuckets are sorted, the distinct buckets within them are not mixed. Figure 5 illustrates the construction steps of a (sorted) superbucket, showing that super-*k*-mers sharing a common minimizer are grouped inside the superbucket. Therefore, a probed *k*-mer is guaranteed to quickly find its bucket, as there is no N character within its (hashed) minimizer. The worst-case scenario, where the first *k*-mer nucleotide is an N, still occurs but within much smaller buckets.

**Figure 5.**
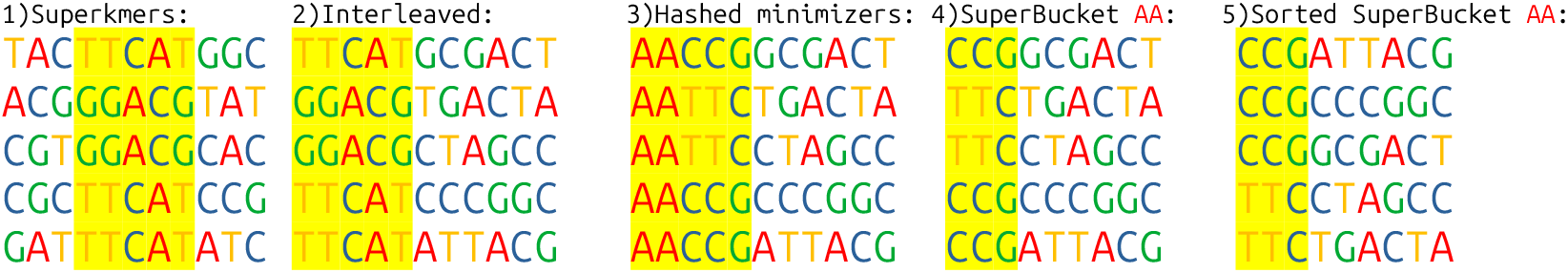
Example showing the successive representation of super-*k*-mers. Minimizers are highlighted in yellow. Super-*k*-mer s are the usual plain text representation of successive *k*-mers sharing a minimizer (1). Interleaved *k*-mers, our introduced transformation, start with the minimizer, with nucleotides closer to the minimizer appearing before those further away (2). Minimizers are then hashed using a reversible hash function to obtain a uniform distribution (3). A superbucket AA is constructed from the first two nucleotides, which are implicit since they all belong to superbucket AA (4). The super-*k*-mers in the superbucket are then sorted to enable fast probing (5).

**Figure 6.**
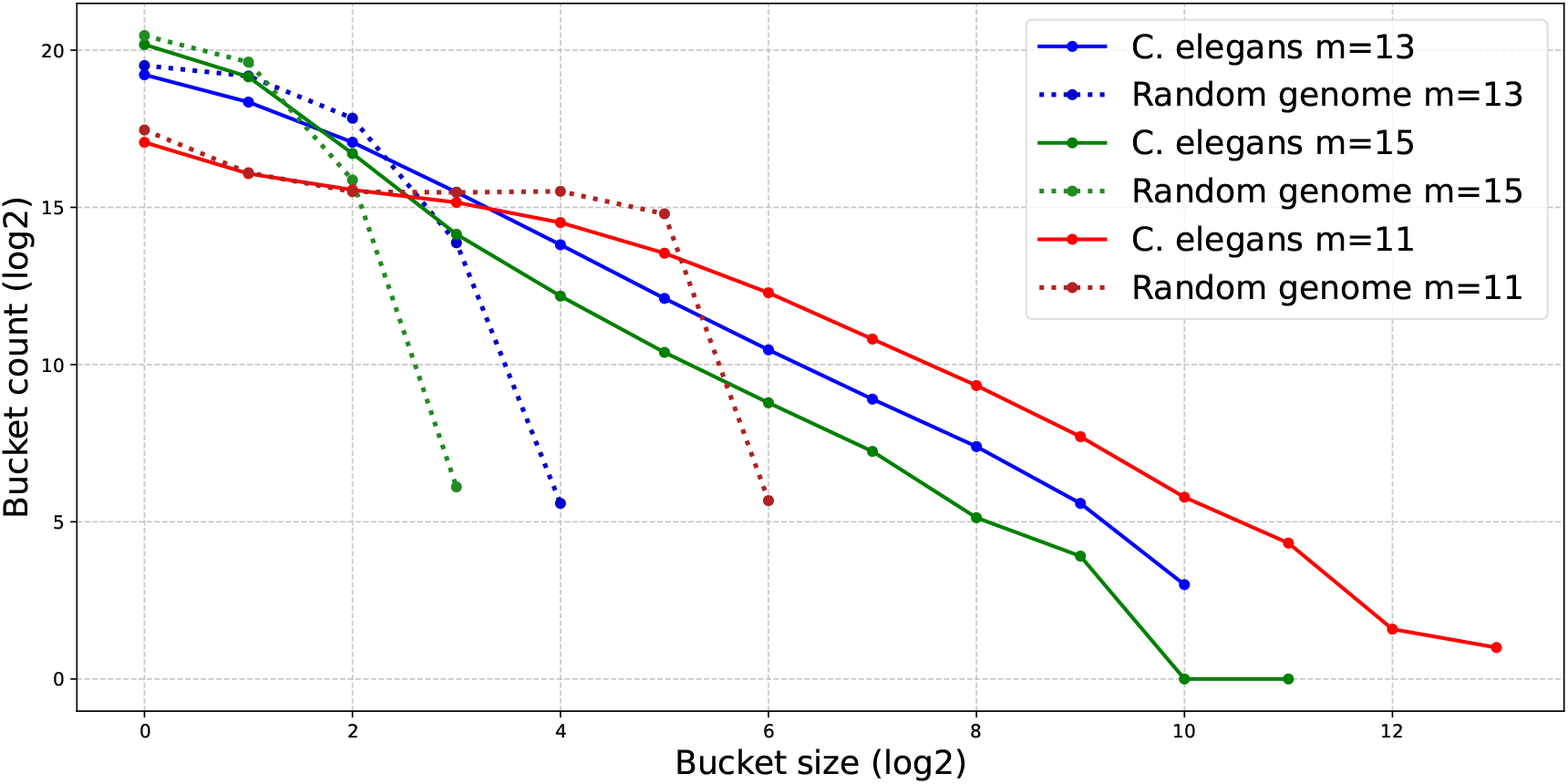
Bucket size distributions comparison between *C. elegans* reference genome and a uniformly distributed random genome of the same size with the same number of buckets (4^11^) but different minimizer sizes: 11, 13 and 15 that correspond to one bucket per minimizer,16 and 256 respectively

### 2.6 Implementation Details

Our goal is to develop a generic dictionary structure utilizing C++ templates, where each *k*-mer is associated with a user-defined data type, such as a 32-bit integer or a 64-bit pointer. In practice, these data are stored in an array alongside the super-*k*-mer list. Specifically, the *i*-th element of the data array corresponds to the *i*-th *k*-mer in the super-*k*-mer list. During sorting operations, the data values are swapped in tandem with their corresponding *k*-mers to maintain the correct associations. To enhance efficiency and avoid the overhead of sorting a large super-*k*-mer list with every insertion, we implement a buffering strategy. A small buffer stores unsorted super-*k*-mers, which are scanned linearly. Once the buffer reaches its capacity, the entire super-*k*-mer list is sorted, and the buffer is subsequently cleared. This approach minimizes the frequency of sorting operations, thereby improving overall performance. The effects of this buffering strategy on performance are further analyzed and discussed in the Results section.

## 3 RESULTS

We implemented the aforementioned strategies in a C++ open-source software named **Brisk** (Brisk Reduced Index for Sequence of K-mers), available at https://github.com/Malfoy/Brisk. Brisk is designed as a C++ library to serve as a dictionary for *k*-mer sequences. Its interface is straightforward and includes the following functionalities:

- **Query Function**: Queries *k*-mers from a sequence and returns a pointer to the associated value of each present *k*-mer.
- **Insert Function**: Inserts *k*-mers from a sequence and returns the associated pointers.
- **Iterator**: Iterates over the indexed *k*-mers and their corresponding values.
- **Serialization Function**: Serializes the data using the KFF format (Dufresne et al., 2022).

As discussed in the introduction, the applications of a *k*-mer dictionary are extensive due to its versatility. To evaluate the performance of Brisk, we focused on one of the simplest and most common uses of such a structure: *k*-mer counting. We compared Brisk against the standard Rust `HashMap`, Jellyfish (Marcais and Kingsford, 2012), a popular *k*-mer counter based on an efficient hash table, and CBL (Martayan et al., 2024a), a *k*-mer set data structure capable of associating a unique identifier to each *k*-mer.

All experiments were conducted on a Supermicro Superserver SYS-2049U-TR4 equipped with 4 Intel SKL 6132 CPUs (14 cores each) @ 2.6 GHz, 3 TB of RAM, running Ubuntu 22.04. To minimize the impact of I/O and parsing operations—which are outside the scope of this work—we aimed to reduce redundancy in the processed documents and optimize the number of distinct *k*-mers. While using random DNA sequences would have been ideal, we opted to evaluate our methods on real-world sequences. Specifically, we focused on bacterial genomes due to their high diversity, which aligns well with our testing requirements. For benchmarking, we downloaded 308,568 bacterial genomes from the NCBI FTP website and randomly selected large subsets of increasing sizes from this collection.

### 3.1 Influence of Parameters

In our first experiment, we benchmarked various bucket sizes to explore the available time-memory trade-off. Figure 7 illustrates the indexing of collections of increasing size using different values of the parameter *b*, which allocates 4^*b*^ distinct buckets. The tested values range from 4^9^ (262k buckets) to 4^17^ (17 billion buckets). For readability, only odd values of *b* are plotted.

**Figure 7.**
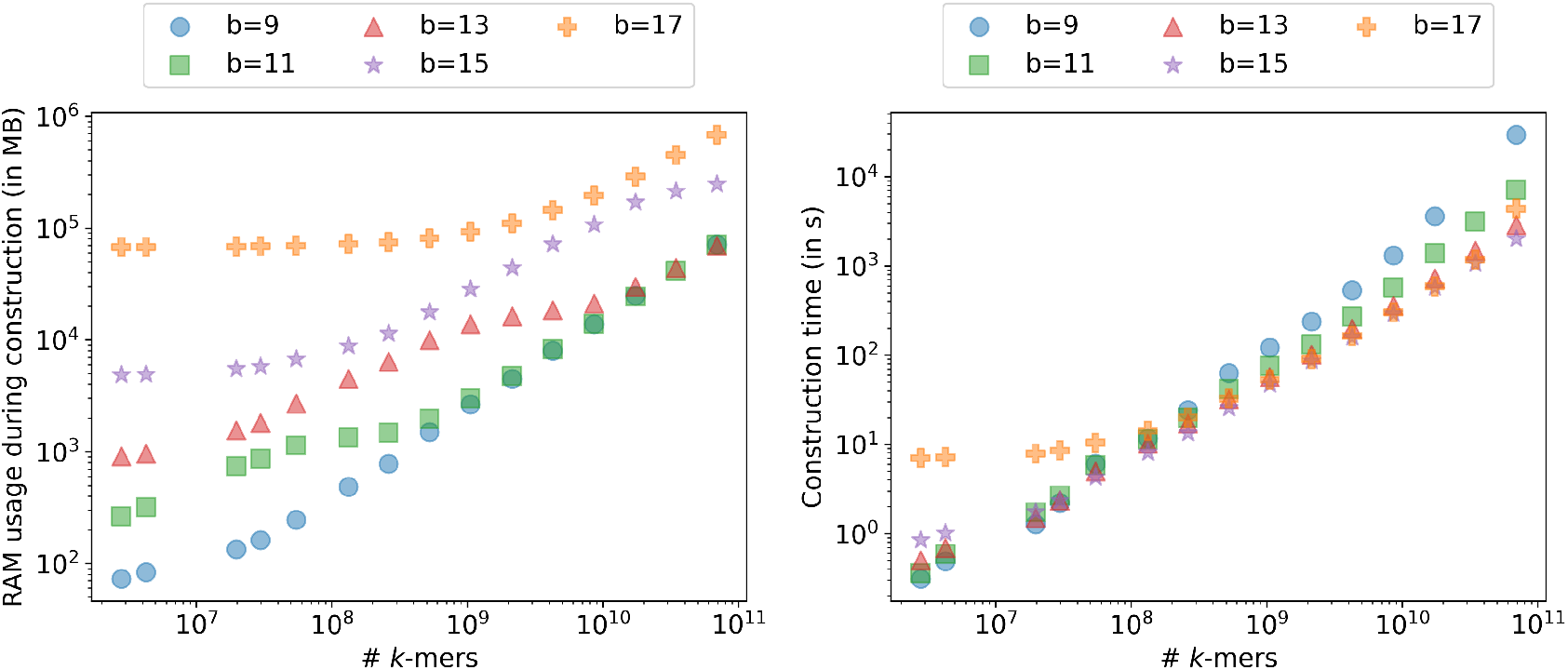
Memory usage (left) and construction time (right) of Brisk with varying values of *b* and *k* = 31, *m* = 21. Higher values of *b* result in increased memory usage but lead to faster construction times.

As expected, the memory overhead of Brisk increases exponentially with *b* for small collections. However, for larger collections, the memory usage becomes dominated by the super-*k*-mers filling the buckets, making it less dependent on *b*. Therefore, larger values of *b* are better suited for larger collections. Additionally, a very small *b* can degrade performance on large collections due to the difficulty in managing very large buckets. Conversely, excessively large *b* values for small collections introduce unnecessary time overhead. Our results indicate that a *b* value around 12 offers a good balance between performance and memory usage across different collection sizes, making it a robust default setting that does not require user expertise.

### 3.2 Multicore Utilization

Brisk’s architecture distributes the dictionary across thousands of substructures, facilitating coarse-grained parallelism. Concurrent access to a given bucket is managed using mutexes to ensure thread safety. In our second experiment, presented in Figure 8b, we measured the wall-clock time for Brisk construction using different numbers of CPU cores.

**Figure 8.**
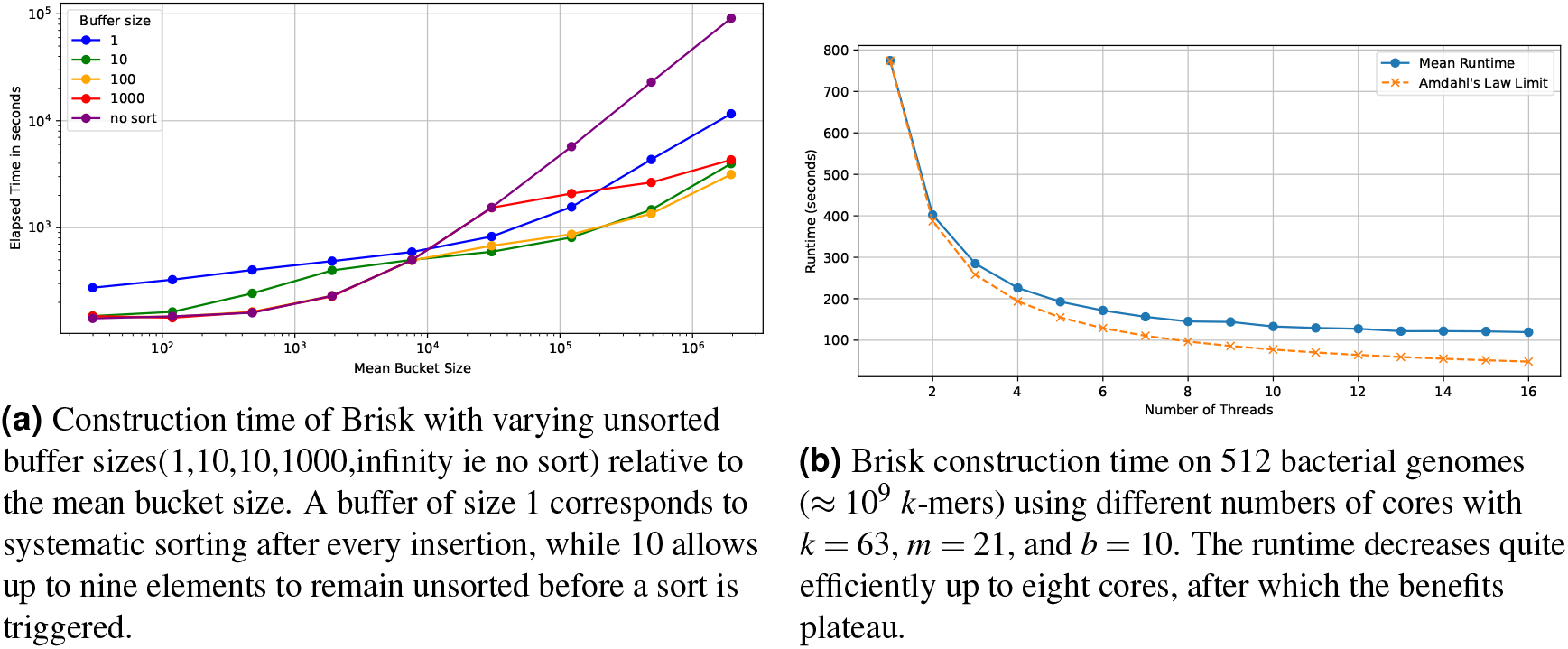
Performance analysis of Brisk construction under varying buffer sizes and multithreading configurations.

The results demonstrate that Brisk scales efficiently with the number of cores up to eight cores, closely approaching the optimal runtime given the fact that some operation like IO can’t be accelerated. Beyond eight CPU cores, the performance gains diminish, likely due to increased contention and management overhead from managing multiple threads.

### 3.3 Sorting Efficiency and Unsorted Buffer Size

In the third experiment, illustrated in Figure 8a, we evaluated Brisk’s performance using different sizes of the unsorted buffer, including a configuration without sorting (*no sort*). Our findings indicate that the novel sorting method employed by Brisk significantly enhances performance. Without sorting, linear probing becomes prohibitively expensive as bucket sizes increase. Conversely, sorted buckets enable more efficient sublinear probing.

Moreover, implementing a buffer to temporarily store unsorted super-*k*-mers reduces the frequency of sorting operations, thereby improving overall performance in practical scenarios.

### 3.4 Comparison with Standard Hashing Methods

We conducted a comprehensive comparison of Brisk, Jellyfish, CBL, and a standard Rust `HashMap` across growing collections of bacterial genomes, ranging from 2^0^ to 2^14^ bacteria. For each dataset, we recorded both the wall-clock time and memory usage. Two *k*-mer sizes were evaluated: *k* = 31, a commonly used value that fits within a 64-bit integer, and *k* = 59, the maximum size supported by CBL.

Figure 9 presents the results for *k* = 31 and *k* = 59. For *k* = 31, Brisk demonstrates significantly lower memory usage compared to the state-of-the-art methods, while matching Jellyfish’s speed. Notably, for *k* = 59, as shown in Figure 9c, the memory consumption advantage of Brisk becomes even more pronounced, and Brisk emerges as the fastest indexing structure.

**Figure 9.**
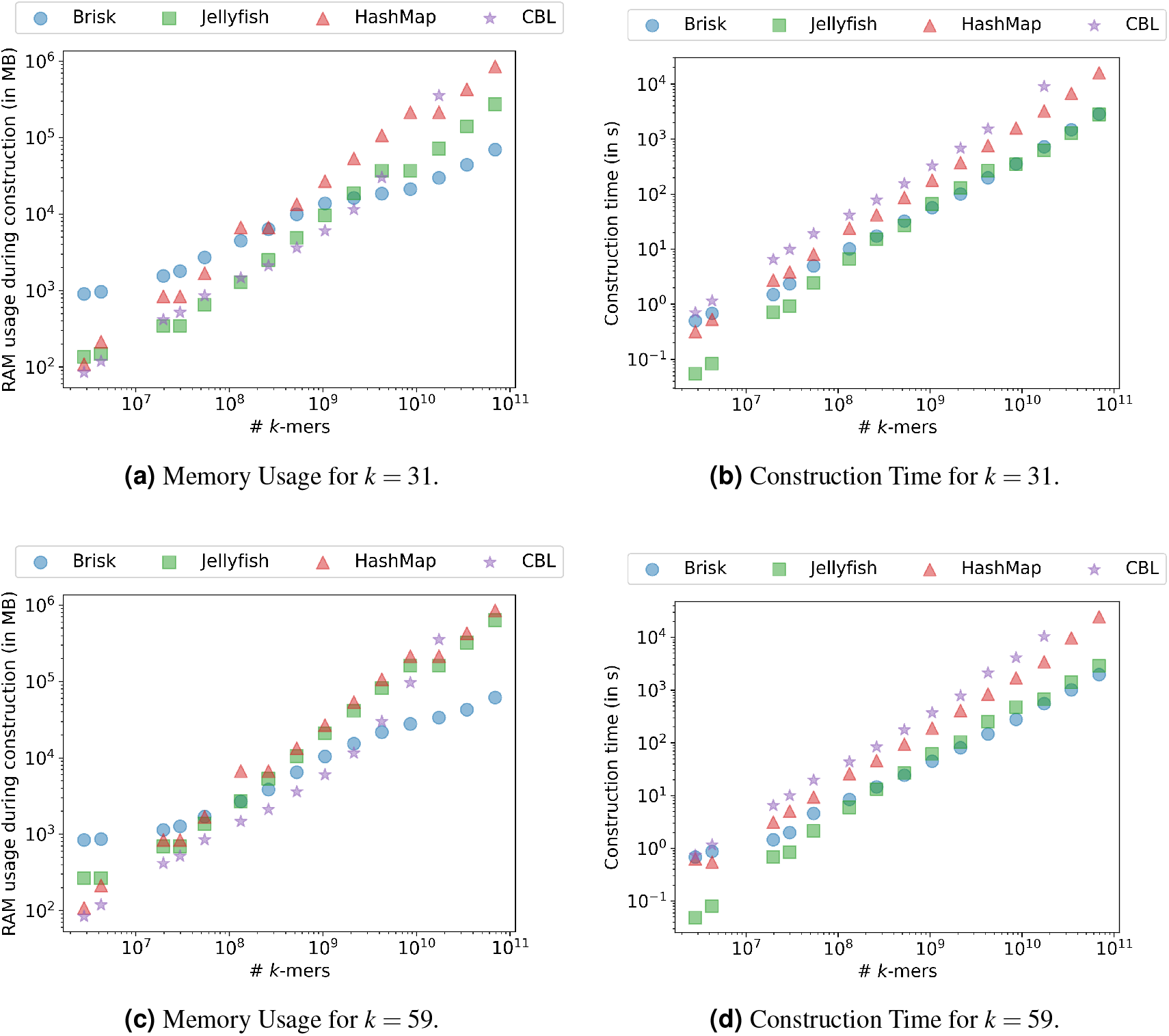
Performance comparison of Brisk, Jellyfish, and Rust `HashMap` across increasing numbers of *k*-mers with *m* = 21, *b* = 13. (a) and (c) depict memory usage, while (b) and (d) show construction times for *k* = 31 and *k* = 59 respectively. Brisk exhibits superior memory efficiency and competitive construction times.

However, it is important to note that the wall-clock time comparisons may not be entirely fair, as CBL and the Rust `HashMap` were not parallelized. To address this, we also performed the same benchmark using a single core, with results displayed in Figure 10. In the single-threaded scenario, Brisk remains comparable to the fast Rust `HashMap` for *k* = 31, albeit slightly slower, and retains its position as the fastest index for *k* = 59.

**Figure 10.**
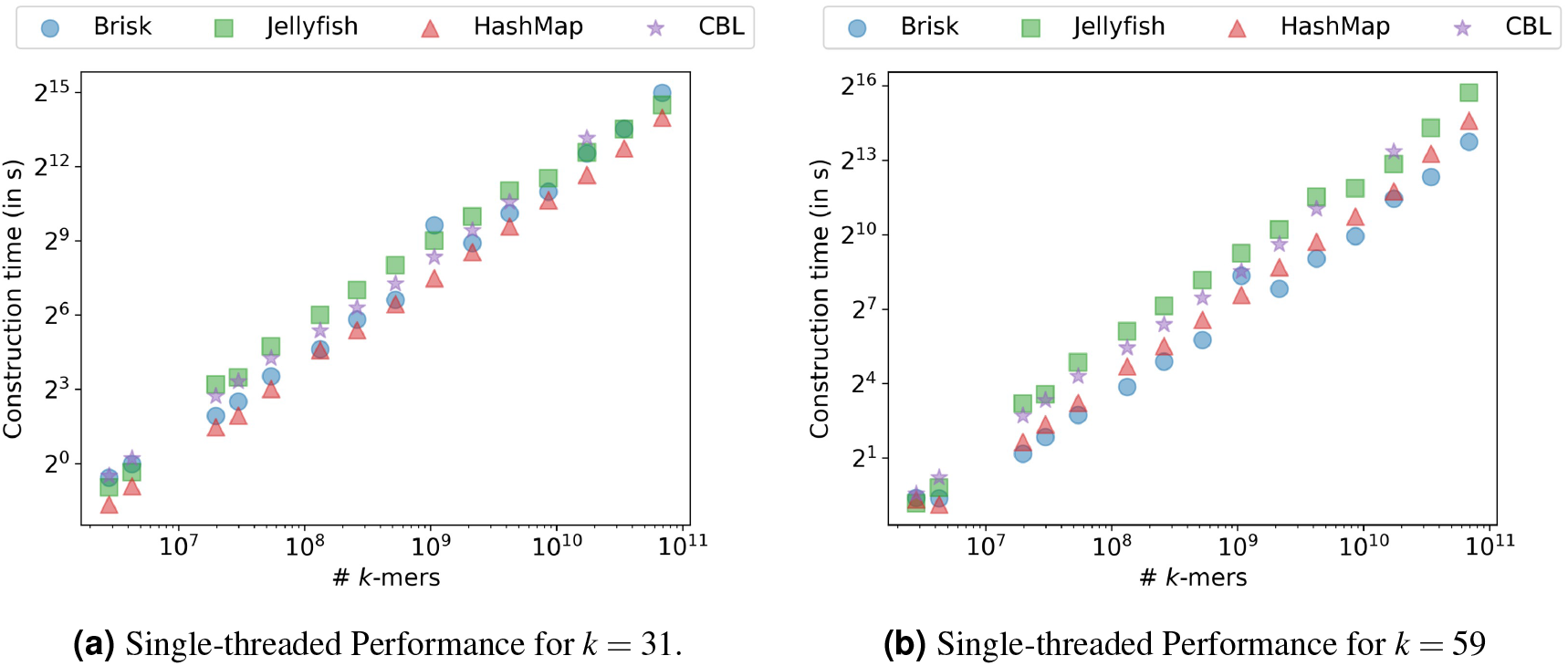
Single-threaded performance of Brisk, Jellyfish, and Rust `HashMap` across increasing numbers of *k*-mers with *m* = 21, *b* = 13. (a) corresponds to *k* = 31, and (b) corresponds to *k* = 59. Brisk maintains competitive performance, especially at larger *k*-mer sizes.

### 3.5 Query Times

In Figure 11, we compare the performance of Brisk in terms of query times for insertions, positive queries (queries for *k*-mers present in the index), and random queries (queries for random *k*-mers, likely absent from the index) as the number of *k*-mers increases. To measure insertion time, an increasing number of bacterial genomes are added to the index. The same dataset is then used to evaluate positive query times. Finally, a FASTA file consisting of random sequences with an equivalent number of *k*-mers is used to assess negative query times. The results show that the throughput of Brisk remains nearly constant as the size of the index grows. Positive queries are several times faster than insertions, while negative queries are even faster, demonstrating an order-of-magnitude improvement over insertion times.

**Figure 11.**
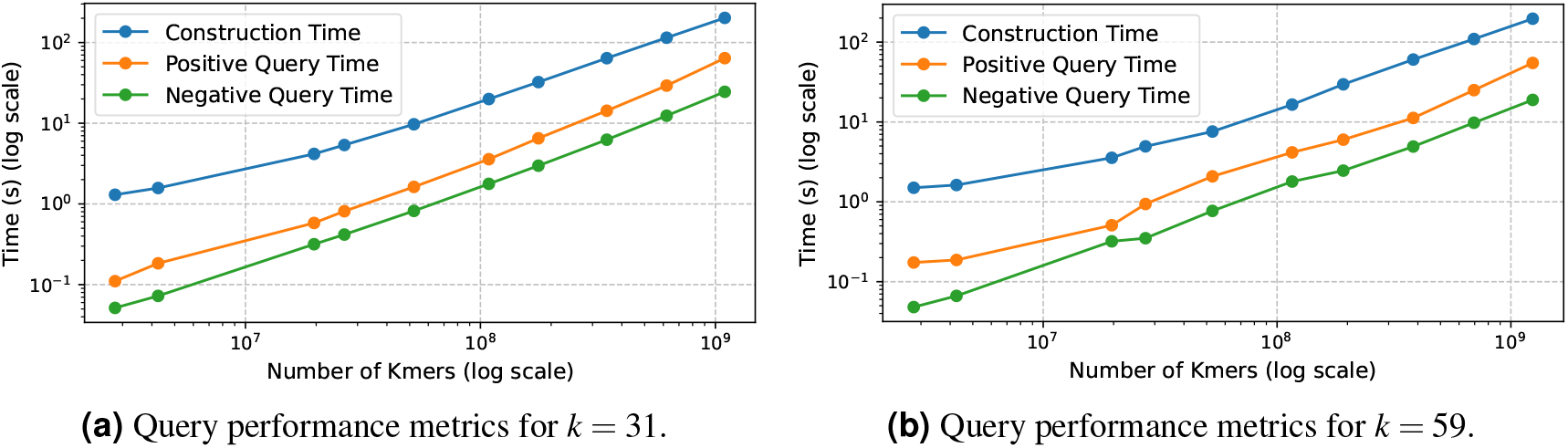
Comparison of positive and random (likely negative) query performances relative to insertion operations across increasing numbers of *k*-mers with *m* = 21, *b* = 13. Sub-figure (a) illustrates results for *k* = 31, while sub-figure (b) shows results for *k* = 59.

## 4 CONCLUSION AND FUTURE WORK

In this study, we introduced a method for adapting a classical dictionary scheme by efficiently indexing super-*k*-mers using a novel interleaved transform. This transform induces a partial ordering, enabling binary search in most cases. Our implementation, Brisk, achieves state-of-the-art throughput while significantly reducing RAM usage. Additionally, Brisk maintains exactness and offers versatility by handling various types of data through templating.

We anticipate that Brisk can serve as a drop-in replacement for traditional *k*-mer dictionaries, thereby enhancing the scalability of numerous tools and workflows. However, there are several features that could be added to further advance this work. For example, extending support to different alphabets, such as proteic or three-dimensional ones, is necessary. Currently, the integer representation of *k*-mers limits the maximum *k*-mer size to 63, as it must be odd and fit within 128-bit integers.

Beyond these implementation-level considerations, the current state of Brisk also raises significant theoretical questions. As it stands, super-*k*-mers are inserted into the index as they are read from the input and remain unchanged thereafter, leaving any empty positions permanently unfilled. Another issue arises when a super-*k*-mer includes a *k*-mer that already exists in the index; in such cases, the new super-*k*-mer cannot be inserted as is, since it would duplicate that *k*-mer within the bucket.

Our current solution is to split the problematic super-*k*-mer into multiple smaller ones to avoid duplication, but this leads to fragmentation that ultimately degrades performance. For the same reason, *k*-mer deletions are not supported, as they would also necessitate splitting. Instead, a *k*-mer can be assigned a “null” value to indicate its removal.

A promising improvement would be to allow new *k*-mers to be inserted directly into existing super-*k*-mers, thereby optimizing memory usage and mitigating fragmentation. However, performing such operations efficiently presents practical challenges. Moreover, it remains theoretically unclear whether a greedy insertion strategy is optimal. This issue is related to the Spectrum Preserving String Set problem but applied to a collection of *k*-mers sharing a minimizer. Another theoretical question, not specific to Brisk, involves finding an improved balance or density in the index and developing more efficient methods for indexing seeds.

Addressing these challenges will not only enhance Brisk’s functionality but also contribute to the broader understanding of *k*-mer indexing strategies.

## 5 COMPETING INTERESTS

No competing interest is declared.

## 6 ACKNOWLEDGMENTS

This work was funded by the French National Research Agency grants AGATE (ANR-21-CE45-0012) and Inception (PIA/ANR16-CONV-0005) and by the research and innovation program under the Marie Skłodowska-Curie grant agreement No 956229 (Alpaca).

## Footnotes

1 https://www.genomeweb.com/sequencing/swedens-single-technologies-progresses-towards-3d-sequencing-10-human-genome

2 https://www.ncbi.nlm.nih.gov/genbank/statistics/

## Notes

https://github.com/Malfoy/Brisk

### Competing Interest Statement

The authors have declared no competing interest.

### Summary of Updates

Most plots have been improved, with enhanced legends and more visible scales. A large number of typos have been corrected. The results sections have been rewritten for clarity and readability.

## REFERENCES

Agret, C., Cazaux, B., and Limasset, A. (2022). Toward Optimal Fingerprint Indexing for Large Scale Genomics. In Boucher, C. and Rahmann, S., editors, 22nd International Workshop on Algorithms in Bioinformatics (WABI 2022), volume 242 of Leibniz International Proceedings in Informatics (LIPIcs), pages 25:1–25:15, Dagstuhl, Germany. Schloss Dagstuhl – Leibniz-Zentrum für Informatik.

Alanko, J., Alipanahi, B., Settle, J., Boucher, C., and Gagie, T. (2021). Buffering updates enables efficient dynamic de bruijn graphs. Computational and structural biotechnology journal, 19:4067–4078.

Alanko, J. N., Vuohtoniemi, J., Mäklin, T., and Puglisi, S. J. (2023). Themisto: a scalable colored k-mer index for sensitive pseudoalignment against hundreds of thousands of bacterial genomes. Bioinformatics, 39(Supplement 1):i260–i269.

Almodaresi, F., Khan, J., Madaminov, S., Ferdman, M., Johnson, R., Pandey, P., and Patro, R. (2022). An incrementally updatable and scalable system for large-scale sequence search using the bentley–saxe transformation.

Almodaresi, F., Sarkar, H., Srivastava, A., and Patro, R. (2018). A space and time-efficient index for the compacted colored de bruijn graph. Bioinformatics, 34(13):i169–i177.

Benoit, G., Raguideau, S., James, R., Phillippy, A. M., Chikhi, R., and Quince, C. (2024). High-quality metagenome assembly from long accurate reads with metamdbg. Nature Biotechnology, pages 1–6.

Bowe, A., Onodera, T., Sadakane, K., and Shibuya, T. (2012). Succinct de bruijn graphs. In International workshop on algorithms in bioinformatics, pages 225–235. Springer.

Chikhi, R., Limasset, A., Jackman, S., Simpson, J. T., and Medvedev, P. (2014). On the representation of de bruijn graphs. In Research in Computational Molecular Biology: 18th Annual International Conference, RECOMB 2014, Pittsburgh, PA, USA, April 2-5, 2014, Proceedings 18, pages 35–55. Springer.

Chikhi, R., Limasset, A., and Medvedev, P. (2016). Compacting de bruijn graphs from sequencing data quickly and in low memory. Bioinformatics, 32(12):i201–i208.

Cracco, A. and Tomescu, A. I. (2023). Extremely fast construction and querying of compacted and colored de bruijn graphs with ggcat. Genome Research, 33(7):1198–1207.

Cremin, C. J., Dash, S., and Huang, X. (2022). Big data: historic advances and emerging trends in biomedical research. Current Research in Biotechnology, 4:138–151.

Deorowicz, S., Kokot, M., Grabowski, S., and Debudaj-Grabysz, A. (2015). Kmc 2: fast and resource-frugal k-mer counting. Bioinformatics, 31(10):1569–1576.

Dufresne, Y., Lemane, T., Marijon, P., Peterlongo, P., Rahman, A., Kokot, M., Medvedev, P., Deorowicz, S., and Chikhi, R. (2022). The k-mer file format: a standardized and compact disk representation of sets of k-mers. Bioinformatics, 38(18):4423–4425.

Ekim, B., Sahlin, K., Medvedev, P., Berger, B., and Chikhi, R. (2023). Efficient mapping of accurate long reads in minimizer space with mapquik. Genome research, 33(7):1188–1197.

Groot Koerkamp, R. and Pibiri, G. E. (2024). The mod-minimizer: A Simple and Efficient Sampling Algorithm for Long k-mers. In Pissis, S. P. and Sung, W.-K., editors, 24th International Workshop on Algorithms in Bioinformatics (WABI 2024), volume 312 of Leibniz International Proceedings in Informatics (LIPIcs), pages 11:1–11:23, Dagstuhl, Germany. Schloss Dagstuhl – Leibniz-Zentrum für Informatik.

Hannoush, K., Marchet, C., and Peterlongo, P. (2024). Cdbgtricks: strategies to update a compacted de bruijn graph. bioRxiv, pages 2024–05.

Holley, G. and Melsted, P. (2020). Bifrost: highly parallel construction and indexing of colored and compacted de bruijn graphs. Genome biology, 21:1–20.

Jenike, K. M., Campos-Domĺnguez, L., Boddé, M., Cerca, J., Hodson, C. N., Schatz, M. C., and Jaron, K. S. (2024). Guide to k-mer approaches for genomics across the tree of life. arXiv preprint arXiv:2404.01519.

Khan, J. and Patro, R. (2021). Cuttlefish: fast, parallel and low-memory compaction of de bruijn graphs from large-scale genome collections. Bioinformatics, 37(Supplement 1):i177–i186.

Li, H. (2018). Minimap2: pairwise alignment for nucleotide sequences. Bioinformatics, 34(18):3094–3100.

Limasset, A., Rizk, G., Chikhi, R., and Peterlongo, P. (2017). Fast and scalable minimal perfect hashing for massive key sets. In 16th International Symposium on Experimental Algorithms (SEA 2017). Schloss Dagstuhl-Leibniz-Zentrum fuer Informatik.

Marcais, G. and Kingsford, C. (2012). Jellyfish: A fast k-mer counter. Tutorialis e Manuais, 1(1-8):1038.

Marchet, C. (2024a). Advancements in colored k-mer sets: essentials for the curious. arXiv preprint arXiv:2409.05214.

Marchet, C. (2024b). Advancements in practical k-mer sets: essentials for the curious. arXiv preprint arXiv:2409.05210.

Marchet, C., Iqbal, Z., Gautheret, D., Salson, M., and Chikhi, R. (2020). Reindeer: efficient indexing of k-mer presence and abundance in sequencing datasets. Bioinformatics, 36(Supplement 1):i177–i185.

Marchet, C., Kerbiriou, M., and Limasset, A. (2021). Blight: efficient exact associative structure for k-mers. Bioinformatics, 37(18):2858–2865.

Marchet, C. and Limasset, A. (2023). Scalable sequence database search using partitioned aggregated bloom comb trees. Bioinformatics, 39(Supplement 1):i252–i259.

Martayan, I., Cazaux, B., Limasset, A., and Marchet, C. (2024a). Conway–Bromage–Lyndon (CBL): an exact, dynamic representation of k-mer sets. Bioinformatics, 40(Supplement 1):i48–i57.

Martayan, I., Robidou, L., Shibuya, Y., and Limasset, A. (2024b). Hyper-k-mers: efficient streaming k-mers representation. bioRxiv.

Marçais, G., Elder, C. S., and Kingsford, C. (2024). k-nonical space: sketching with reverse complements. Bioinformatics, 40(11):btae629.

Pellow, D., Pu, L., Ekim, B., Kotlar, L., Berger, B., Shamir, R., and Orenstein, Y. (2023). Efficient minimizer orders for large values of k using minimum decycling sets. Genome Research, 33(7):1154–1161.

Pibiri, G. E. (2022). Sparse and skew hashing of k-mers. Bioinformatics, 38(Supplement 1):i185–i194.

Pibiri, G. E., Shibuya, Y., and Limasset, A. (2023). Locality-preserving minimal perfect hashing of k-mers. Bioinformatics, 39(Supplement 1):i534–i543.

Pibiri, G. E. and Trani, R. (2021). Pthash: Revisiting fch minimal perfect hashing. In Proceedings of the 44th international ACM SIGIR conference on research and development in information retrieval, pages 1339–1348.

Rahman, A. and Medevedev, P. (2021). Representation of k-mer sets using spectrum-preserving string sets. Journal of Computational Biology, 28(4):381–394.

Rouzé, T., Martayan, I., Marchet, C., and Limasset, A. (2023). Fractional Hitting Sets for Efficient and Lightweight Genomic Data Sketching. In Belazzougui, D. and Ouangraoua, A., editors, 23rd International Workshop on Algorithms in Bioinformatics (WABI 2023), volume 273 of Leibniz International Proceedings in Informatics (LIPIcs), pages 15:1–15:27, Dagstuhl, Germany. Schloss Dagstuhl – Leibniz-Zentrum für Informatik.

Schleimer, S., Wilkerson, D. S., and Aiken, A. (2003). Winnowing: local algorithms for document fingerprinting. In Proceedings of the 2003 ACM SIGMOD international conference on Management of data, SIGMOD ‘03, pages 76–85, New York, NY, USA. Association for Computing Machinery.

Schmidt, S. and Alanko, J. N. (2023). Eulertigs: minimum plain text representation of k-mer sets without repetitions in linear time. Algorithms for Molecular Biology, 18(1):5.

Schmidt, S., Khan, S., Alanko, J. N., Pibiri, G. E., and Tomescu, A. I. (2023). Matchtigs: minimum plain text representation of k-mer sets. Genome Biology, 24(1):136.

Sladky, O., Vesely, P., and Břinda, K. (2023). Masked superstrings as a unified framework for textual k-mer set representations. bioRxiv, pages 2023–02.

Stephens, Z. D., Lee, S. Y., Faghri, F., Campbell, R. H., Zhai, C., Efron, M. J., Iyer, R., Schatz, M. C., Sinha, S., and Robinson, G. E. (2015). Big data: astronomical or genomical? PLoS biology, 13(7):e1002195.

Zheng, H., Kingsford, C., and Marçais, G. (2020). Improved design and analysis of practical minimizers. Bioinformatics, 36(Supplement 1):i119–i127.

